# Nonsense correlations in neuroscience

**DOI:** 10.1101/2020.11.29.402719

**Authors:** Kenneth D. Harris

## Abstract

Many neurophysiological signals exhibit slow continuous trends over time. Because standard correlation analyses assume that all samples are independent, they can yield apparently significant “nonsense correlations” even for signals that are completely unrelated. Here we compare the performance of several methods for assessing correlations between timeseries, using simulated slowly drifting signals with and without genuine correlations. The best performance was obtained from a “pseudosession method”, which relies on one of the signals being randomly generated by the experimenter, or a “session perturbation” method which requires multiple recordings under the same conditions. If neither of these is applicable, a “linear shift” method can be used when one of the signals is stationary. Methods based on cross-validation, circular shifting, phase randomization, or detrending gave up to 100% false positive rates in our simulations. We conclude that analysis of neural timeseries is best performed when stationarity and randomization is built into the experimental design.

---

In neuroscience we often aim to find correlations between variables that depend on time. For example, we might correlate neuronal population activity on each trial of a task with behavioral variables such as choices. The statistical analysis of such data is difficult because the recorded variables often show slow changes in activity, which can lead to apparent correlations between them even if they are completely unrelated.

This phenomenon was given the memorable name “nonsense correlation” by statistician G. Udny Yule (Yule, 1926). Yule illustrated this problem with an apparent correlation between mortality rates and the fraction of marriages conducted by the Church of England. A more recent illustration describes an apparent correlation between cryptocurrency prices and activity in the brains of mice (Meijer, 2021).

The problem of nonsense correlations has been discussed extensively in fields such as econometrics (Box, 2008; Granger and Newbold, 1974; Haugh, 1976; Phillips, 1986), but despite its importance to understanding neurophysiology data, has seen little discussion in this field (but see Elber-Dorozko and Loewenstein, 2018).

Here, we evaluate ten possible solutions to the problem, by applying them to simulated neural data. Two methods (the pseudosession and session permutation methods) do not produce nonsense correlations, however they cannot be used in all situations. The conservative linear shift method does not produce nonsense correlations if one of the series is stationary. The remaining methods (naïve correlation, circular shift, phase and wavelet randomization, cross-validation, auto-decorrelation) can all produce nonsense correlations. We end with suggestions for how to design experiments on which pseudosession and session permutation methods can be used.

## What are nonsense correlations?

To illustrate the phenomenon of nonsense correlations, we consider a simulated experiment (**Figure 1**). Imagine we have recorded a population of *C* = 10 cells and computed their firing rate on *T* = 200 behavioral trials. To simulate the case that the neurons encode no information about behavior, we generate their rates randomly, independent of each other and of the simulated behavioral variables. We simulate slow rate drifts by summing logistic sigmoid functions centered on random times, together with pink noise (Methods; **Figure 1A1,1A2**).

**Figure 1.**
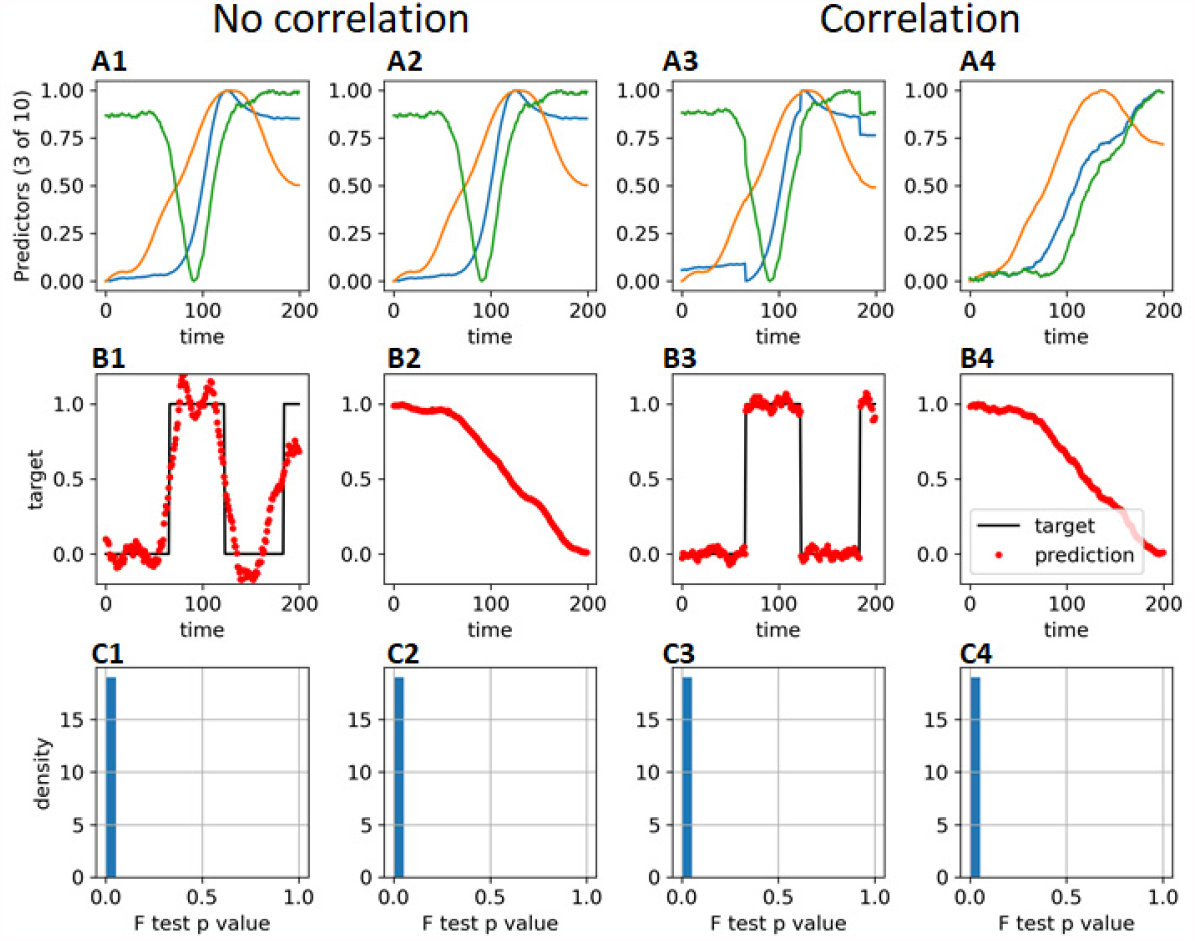
Naïve correlation method. **A1-A4:** firing rates of 3 out of 10 simulated neurons as a function of trial number. In columns 1 and 2 the simulated firing rates are uncorrelated with the behavioral variable. In columns 3 and 4, a small randomly weighted copy of the behavioral variable has been added to the firing rates to produce a correlation. **B1-B4:** behavioral variable (black) and prediction of it from neural firing (red dots), using multiple linear regression with weights constant across the simulation. **C1-C4:** histogram of F-test p-values measuring significance of the linear regression over 1000 simulations.

We consider two types of simulated behavioral variable. First, we consider a binary “block” variable, which switches pseudo-randomly during the experiment; for example, this could indicate which of two stimuli or actions is most likely to be rewarded (**Figure 1B1**). Even though the block variable was generated independently of the neural activity, it is possible to predict it accurately from neural activity, since by chance some of the neurons showed rate shifts at times close to the block switches (e.g. the green cell in **Figure 1A1**).

The second type of simulated behavioral variable was a continuous one, simulated the same way as the neural variables (**Figure 1B2)**; for example this could measure running speed on each trial. Again, this variable could be predicted almost perfectly from the simulated neural activity, even though it was generated independently.

To simulate a situation where neuronal firing rates do encode information about the behavioral variable, we added a small multiple of the behavioral variable to each neuron’s firing rate, with a random weight (**Figure 1A3, 1A4**). Throughout the paper we consider these four scenarios: a binary or continuous behavioral variable, with or without genuine correlation to neural activity.

To measure how well the behavioral variable correlated with neural activity on each trial, we predicted it by multiple linear regression. (Other methods could be used but this would not change the basic results.) Naively applying the usual test of significance for multiple linear regression (the F-test), we find statistical significance in each of our four scenarios, for each of 1000 simulations (**Figure 1C**).

Naïve correlation thus always produced a false-positive error even when there was no genuine relationship between neural activity and the behavioral variable. This is because the F test assumes that the data on each timestep are statistically independent. However, both the predictor firing rates, and the target behavioral variable are correlated across timesteps, and the test gives false significance.

## Defining correlation between time series

Before considering potential solutions to the problem of nonsense correlations, we must first clearly define what we mean by a correlation between time series. To do so, we recall some basic concepts of probability theory, working here within the classical “frequentist” framework.

A fundamental concept in probability theory is the *sample space*. The sample space defines the set of all possible outcomes of an experiment, and a point in the sample space determines the value of everything measured in the experimental session. Throughout this paper we consider a simultaneous recording of *C* cells and one behavioral variable, both measured on *T* trials. A point in the sample space is therefore defined by (*C* + 1)*T* numbers: the firing rate of each neuron and the behavioral variable on each trial. We will denote the firing rate of cell *c* on trial *t* as *x*_*ct*_, and gather them together into an *CT*-dimensional vector **x;** and we will denote the behavioral variable on trial *t* as *y*_*t*_, gathered together into a *T*-dimensional vector **y**. Importantly, the sample space is defined by the entire history of these variables on all trials, not by their values on a single trial.

In the frequentist framework, we consider experiments to be repeatable, at least in principle: even if we only performed the experiment once, we consider it as part of an ensemble of repeats we could have performed. A probability distribution ℙ(**x, y**) measures the frequency with which the experiment yields a particular outcome, over the infinite ensemble of possible repetitions of the experiment.

We say that neural activity is uncorrelated with behavior if the entire history of neural activity in an experiment (summarized by the vector **x**) is statistically independent of the history of behavioral variables summarized in **y**, i.e. if ℙ(**x, y**) = ℙ(**x**)ℙ(**y**).

Importantly, this definition allows neural activity to be *autocorrelated*: the firing rate of neuron *n* on trial *t* can be correlated with the firing rate of neuron *m* on trial *u*. Behavior can also be autocorrelated: the value of the behavioral variable at one time can be correlated with the value at another. Instead, independence requires that there be no *cross-correlation:* the activity of any neuron at any time is independent of behavior at any time. Thus, ℙ(*x*_*ct*_, *y*_*u*_) = ℙ(*x*_*ct*_)ℙ(*y*_*u*_), for any cell *c* and any pair of times *t* and *u*.

A correlation between timeseries is therefore defined as a relationship that holds consistently across multiple repeats of the experiment, rather than across timepoints within a single experimental session. Predicting behavior from neural activity within a single session (Figure 1) does not show that neural activity is correlated with behavior. Instead, it must be possible to predict behavior from activity for all experimental sessions, using the same set of prediction weights for each session.

Does this mean that to show a correlation between neural activity and behavior one must record from the same neural population over multiple experimental sessions? Luckily, the answer is no, provided we make certain further assumptions. We next discuss how different assumptions allow different methods for detecting true correlations between time series. We focus on the simple case of testing whether neural and behavioral variables are correlated: more complex questions such as testing whether neural activity correlates with some behavioral variables after taking others into account, are discussed elsewhere (Harris, 2021) and summarized briefly at the end of this manuscript.

## Pseudosession method

The “pseudosession method” is simple, requires only a single experimental session, and is the only method we describe here that can show a causal relationship between two timeseries. However it has the strongest requirement: that one of the timeseries is randomly generated by the experimenter according to a known probability distribution. This method could be used for example to test whether neural activity differs between behavioral blocks, in an experiment where the block structure is generated randomly without dependence on the subject’s choices.

Let **x** and **y** denote the histories of neural activity and of the behavioral variable in a single session. The pseudosession method requires a user-specified function *V*(**x, y**), a single real number which quantifies the degree of association between **x** and **y** in that session. A good choice is the error of a classifier trained to predict **y** from **x** or vice versa. This error can be cross-validated but does not need to be. In fact, any choice of *V* gives a valid test; poor choices can only result in false-negative errors. In this paper we use the squared error of linear regression summed over time points. But any classifier can be used, including if it uses multiple timepoints of one series to predict individual timepoints of the other.

To apply the pseudosession method, we repeatedly generate random histories **y**′_*i*_ from the same probability distribution that generated **y**, refit the prediction model to predict each 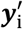 from **x**, and recompute the test statistic *V*(**x, y**′_*i*_). We then define a p-value as the quantile of *V*(**x, y**) relative to the null ensemble {*V*(**x, y**′_*i*_)}. This method therefore rejects the null hypothesis of no correlation if we can predict the actual behavioral history significantly better than we could predict a randomly generated one.

Applying the method to our four scenarios (**Figure 2**), we observe that p-values are evenly distributed when there is no true correlation but concentrated near zero when there is. We conclude that the pseudosession method works reliably when the behavioral variable is generated randomly from a known distribution.

**Figure 2.**
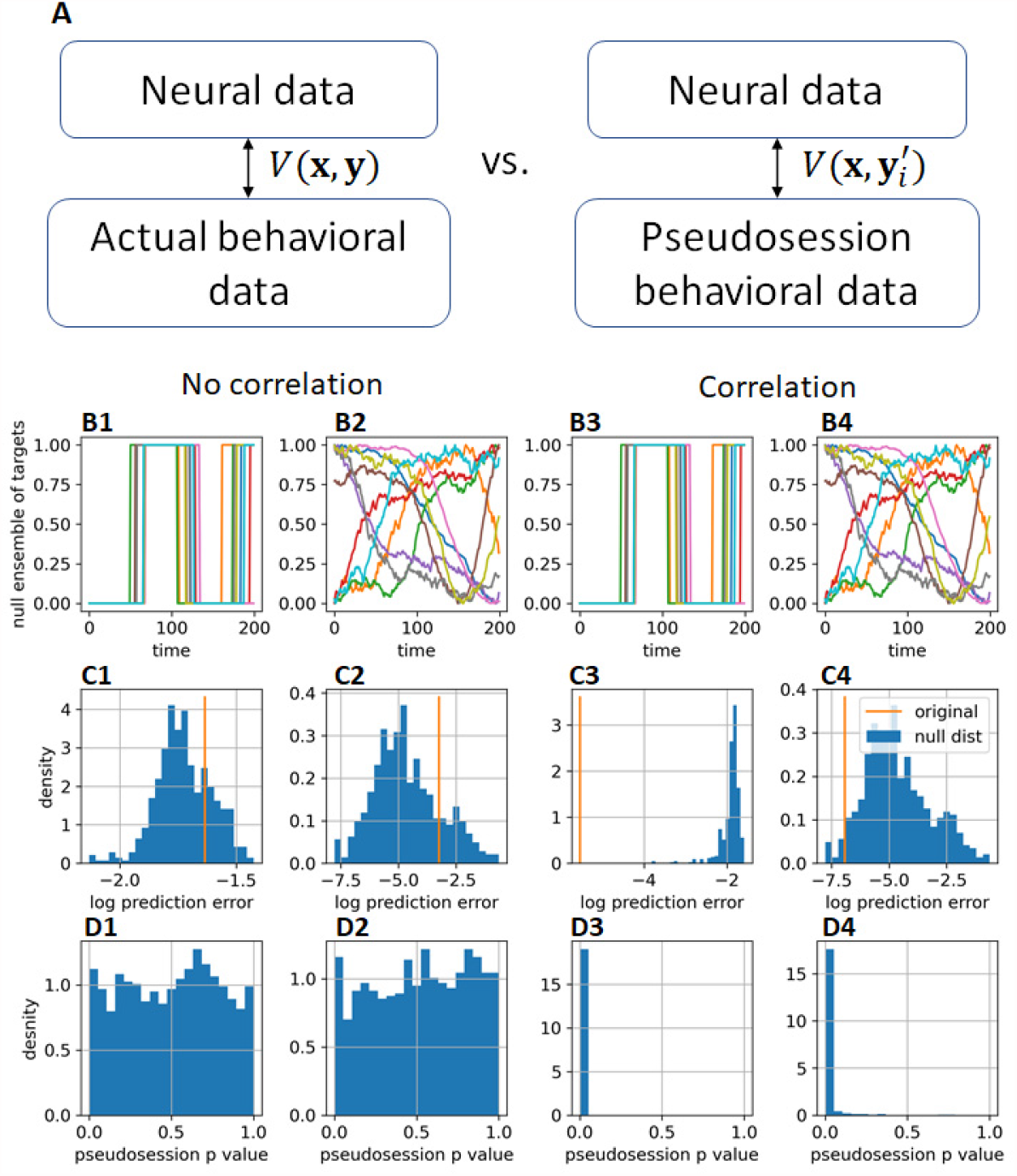
Pseudosession method. **A**: a test statistic *V*(**x, y**) is computed that measures the strength of relationship between the neural and behavioral data for an experiment. This is compared against a null distribution of 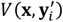 obtained by repeatedly generating other behavioral data 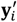 drawn from the same distribution as **y. B:** ensemble of behavioral variables for 10 pseudosessions, drawn from the same probability distribution as the original. **C**: histogram of log prediction error 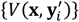 of null distribution (blue), with *V*(**x, y**), the value for original data superimposed (orange). **D:** histogram of p-values obtained by comparing the predictability of the actual behavioral variable against a null ensemble of predictability of pseudosessions, from 1000 simulations. Columns 1-4 correspond to the same scenarios as in Figure 1.

## Session permutation method

Because the pseudosession method requires the behavioral variable to have been randomly generated by the experimenter, it cannot be used to correlate neural activity with variables such as the subject’s choices or running speed, which are not under the experimenter’s control. The session permutation method can analyze these cases but requires data from multiple sessions recorded under identical conditions.

The session permutation method asks whether neural activity predicts the behavioral variable on the same session more accurately than on other sessions. We denote the vectors containing the history neural activity and behavioral variables on the *s*^*th*^ session as **x**_*s*_ and **y**_*s*_. We sum the association measure over sessions to obtain a test statistic 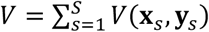. We compare this test statistic to a null ensemble in which the neural data of each session is compared to behavioral data from a randomly chosen session: 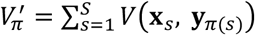, where *π* runs over all of the *S*! permutations of the *S* sessions. To obtain statistical significance needs at least 5 sessions (since 5! = 120).

This method works for all our four scenarios, giving a flat distribution of p-values when the null hypothesis is true, and a sharp peak near 0 when the null is false (**Figure 3)**.

**Figure 3.**
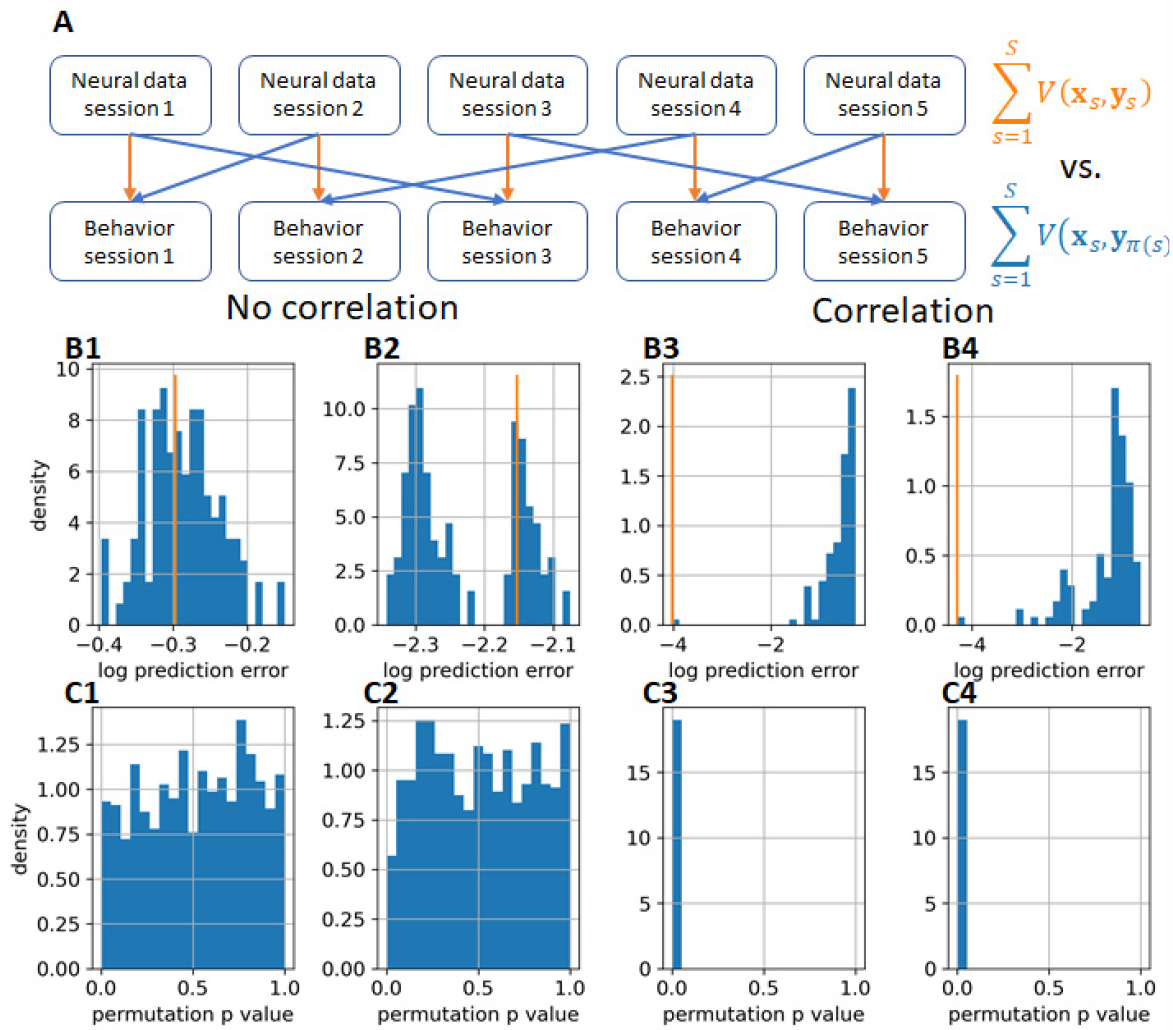
Session permutation method. **A:** a test statistic is computed by summing the predictability of behavioral from neural data over each of *S* total sessions. This is compared to a null ensemble obtained from each of the *S*! possible permutations of the sessions. **B:** histogram of log prediction error of permuted sessions (blue), with value for unpermuted data superimposed (orange). **C:** Histogram of p-values obtained with this method, over 1000 simulations. Columns 1-4 correspond to the same scenarios as in Figure 1.

Session permutation does not require that the same neurons be recorded in each session, provided one can consider the experiments to be statistically independent. The statistic *V*(**x**_*s*_, **y**_*s*_) measures the degree to which the recorded population predicts behavior, and this can be computed using different neural populations on different sessions. If one cannot return to the same neurons on each session, however, it is not possible to say which neurons correlate with the behavioral variables; it is only possible to conclude that the population as a whole does.

Some caution is required in interpreting results of the session permutation method. Without a randomized experimental design, one cannot infer causality as there may be a third factor affecting the neural and behavioral recordings. For example, if the *S* sessions were recorded sequentially from the same subject, and consecutive experiments showed both a degradation in both the quality of neuronal recording and in behavioral performance, one might observe a correlation between neural activity and behavior simply for this reason. If some sessions were recorded from different subjects one might observe a correlation between neural activity and behavior, simply because the subjects used different behavioral strategies and their recordings contained different numbers of neurons.

## Linear shift method

We have defined correlation between neural activity and behavior as a relationship holding consistently across sessions. Nevertheless, we can infer a correlation from just one session, if we make further assumptions. The linear shift method does this by assuming *stationarity*.

A probability distribution for time series is said to be stationary if it is invariant to time shifting: for any 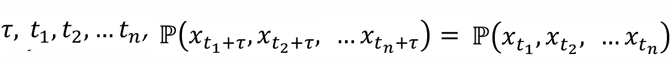. Stationarity is a property of the ensemble of all possible histories, rather than of any one session. Stationarity does not mean that the data have a temporally consistent character during a single experiment; rather, it means that absolute time is irrelevant to the ensemble. For example, our ensemble of simulated block histories (**Figure 2B1**) is not stationary, since the first trial of any session is always in block 0; it could be made stationary by first generating a long block history, then starting at a random point. Experimentally measured data such as behavioral and neural timeseries will be nonstationary if they show consistent trends from the beginning to end of all experiments. For example, if subjects typically respond faster at the beginning of a session experiment than at the end, then the timeseries of reaction times would be nonstationary.

The linear shift method (Harris, 2020) tests the null hypothesis that two time series are independent, and one of them is stationary (in this case we assume **x**). It is based on the idea that if the series are correlated, it should be easier to predict a segment of one them from a simultaneous segment of the other, than from a temporally shifted segment.

In detail, let **x**[*a*: *b*] denote the segment of **x** from trial *a* to trial b−1. Given an integer parameter 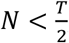, the linear shift method calculates the association of a central segment of **y** with a shifted segment of **x**. For all *s* = −*N* … *N*, it computes *V*_*s*_ = *V*(**x**[*N* + *s*: *T* − *N* + *s*], **y**[*N*: *T* − *N*]), where *V* is again a measure of association between neural activity and behavior such as the error of predicting behavior from neural activity. A test statistic 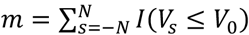 counts how many shifts produce an association as least as strong as the unshifted data (*I* is an indicator function: 1 if its argument is true, 0 if false). We reject the null hypothesis at significance level *α* if *m* ≤ *α*(*N* + 1). This is a conservative test: for any function *V*, the probability that a valid null will be rejected is guaranteed to be less than *α* if **x** and **y** are independent and **x** is stationary. An approximate test rejecting the null if *m* ≤ *p*(2*N* + 1) becomes approximately valid in the limit *N* → ∞, but for any finite *N* there exist (increasingly strange) counterexamples for which the approximate test rejects a valid null with probability more than *α* (Harris, 2020). Note that previous version of the present manuscript suggested using an initial rather than central segment of **y**, but that approach does not allow a conservative test and is not recommended.

We evaluated the linear shift method on the same four scenarios as before (**Figure 4**). We applied the test with *N* = 19, for which the conservative test will reject the null at p=0.05 if *m* = 1, meaning the unshifted central segments of **x** and **y** are more strongly associated than any of the shifted segments, and the approximate test will reject the null at p=0.051 if *m* ≤ 2, meaning that at most one shifted value can have a stronger association. For uncorrelated data, the conservative test falsely rejected the null in 2.7% and 2.9% of simulations of the block and continuous **y**; the approximate test rejected in 4.2% and 5.1% of simulations. When a genuine correlation was present, both tests rejected the null in 100% of block simulations; for continuous simulations, rejection rates were 89% and 91%. Thus, the approximate test did not produce excess false positive errors in these simulations, although it provided little advantage. We also evaluated the test shifting **y** rather than **x**, which in the block scenario is not stationary. Nevertheless, the conservative and approximate tests rejected a valid null in just 1.9% and 3.0% of simulations, indicating that non-stationarity did not yield false-positive errors in this case.

**Figure 4.**
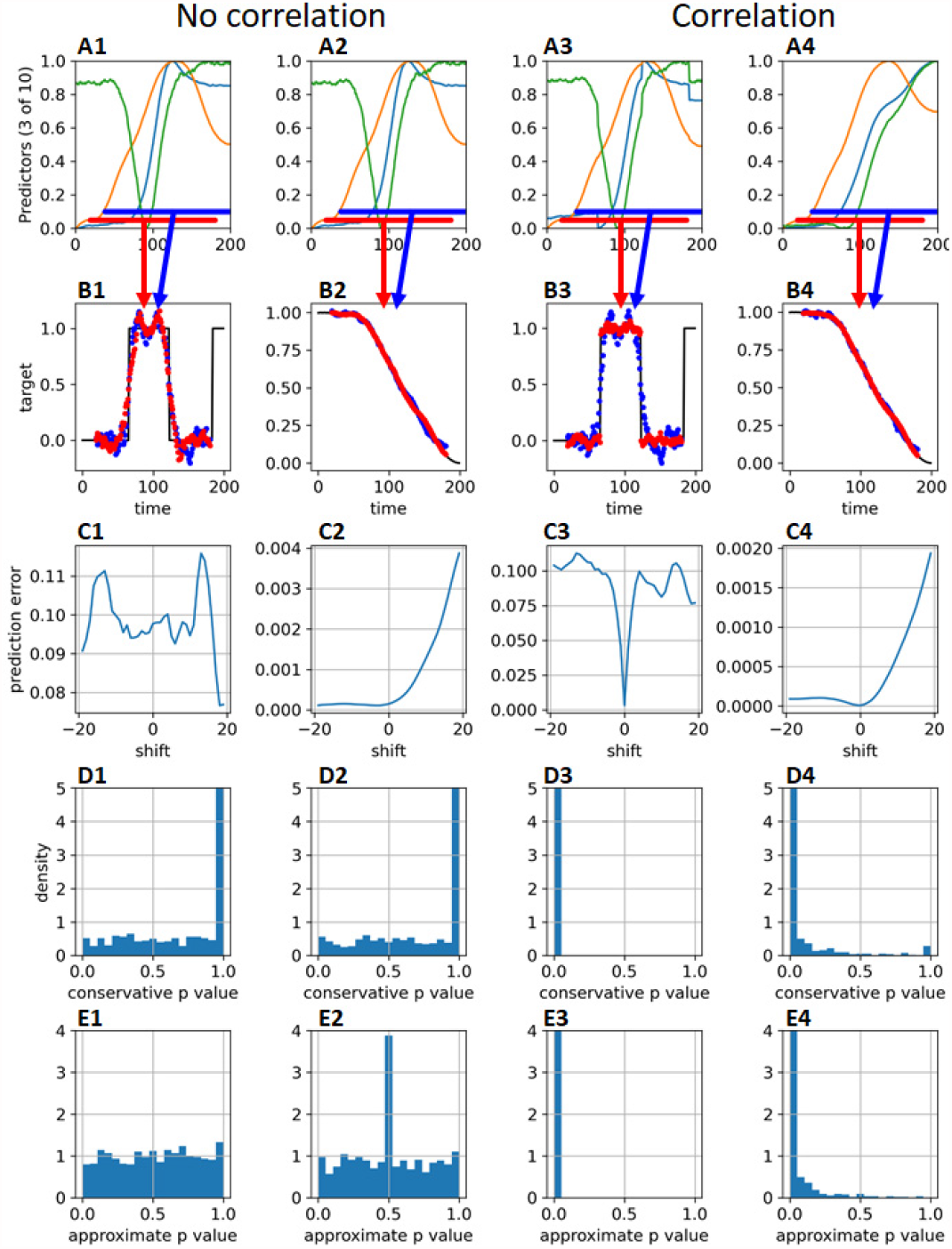
Linear shift method. **A:** Neural data as in Figure 1. Red bar indicates time segment used to make unshifted prediction; blue shows one example shift. **B:** the central segment of the behavioral data is predicted from unshifted (red) or shifted (blue) neural data. For correlated series, the unshifted predictor is more accurate (B3, B4). **C:** Prediction error as a function of shift. **D,E:** histogram of p-values obtained using the conservative or approximate methods, over 1000 simulations. The peak at p=0.5 in panel E2 reflects cases where the prediction error depends monotonically on shift. The four columns refer to the four scenarios of Figure 1.

Thus, the conservative linear shift test can be safely used when one has reason to believe that one of the time series being compared is stationary. Neither the approximate method nor nonstationarity produced excess false rejections in our simulations, but this does not guarantee safety of those methods in all situations, and the twofold loss in statistical power of the conservative method is not a high price to pay for the increased safety it provides from false positive errors.

## Circular shift method

An alternative to the linear shift method is to generate a null ensemble by circularly shifting one of the timeseries: to replace **x**[0: *T*] with the concatenation of **x**[*s*: *T*] and **x**[0: *s*]. This has the advantage of using all the data, unlike the linear shift method which discards some. However, in our simulations circular shifting showed much greater inflation of false-positive errors than linear shifting (**Figure 5**).

**Figure 5.**
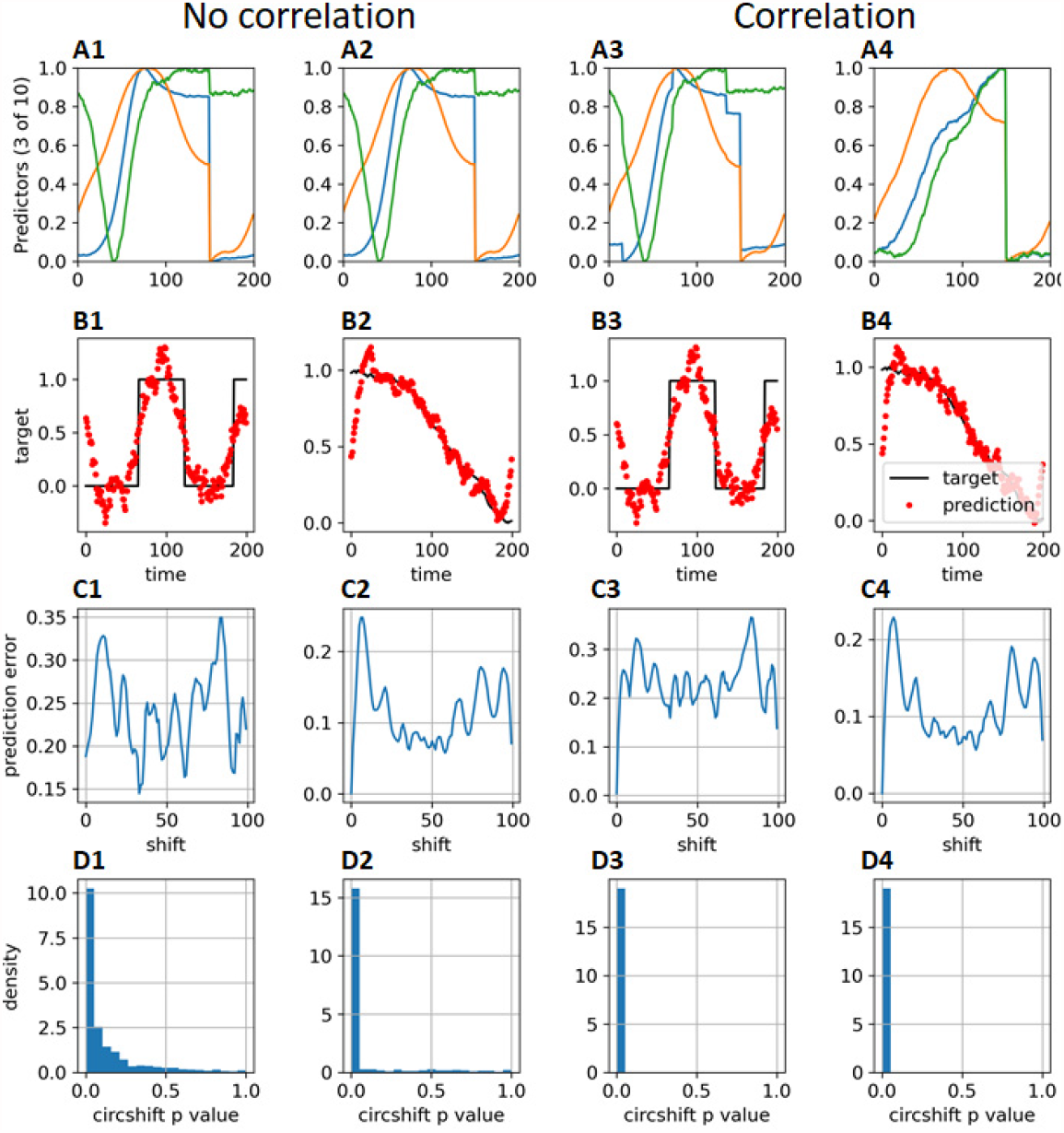
Circular shift method. **A,B:** To create a null distribution, the neural data is circularly time shifted and used to predict behavioral data. The example shows a circular shift of 75 trials, where a discontinuity is visible. **C:** Prediction error as a function of circular shift magnitude. **D:** histogram of p-values obtained for the method, over 1000 simulations. Columns refer to the four scenarios of Figure 1.

The reason for this problem is that the circular shift method makes an assumption that is unlikely to hold. It tests the null hypothesis that not only are the series independent, but one of them is also cyclo-stationary: i.e. the probability of observing a particular history is the same as the probability of observing a cyclic shift of it. The reason this is unlikely to hold is that unless the start and end values of the timeseries are identical, cyclic shifting will introduce a discontinuity (**Figure 5A**), which then renders the prediction of the behavioral series worse.

## Phase/wavelet randomization

Another alternative to linear shifting, which has been suggested in the fMRI literature (Bullmore et al., 2001; Laird et al., 2004) is phase or wavelet randomization.

In the phase randomization method, a null distribution is obtained by applying a Fourier transform to one of the timeseries, multiplying each Fourier coefficient by a random phase and reverse transforming.

Our simulations suggested that this method inflates false-positive errors much more than the linear shift method (**Figure 6**). Whereas the circular shift method transforms continuous timeseries into discontinuous ones, phase randomization instead imposes cyclic continuity: after phase randomization the last sample is always close to the first. Furthermore, the phase-randomized signals have more high-frequency activity than the original, as the high-frequency energy resulting from the cyclic discontinuity in the original data has now been spread throughout time. As a result, the phase randomized data tends to predict the behavioral data worse, resulting in inflated false-positive errors.

**Figure 6.**
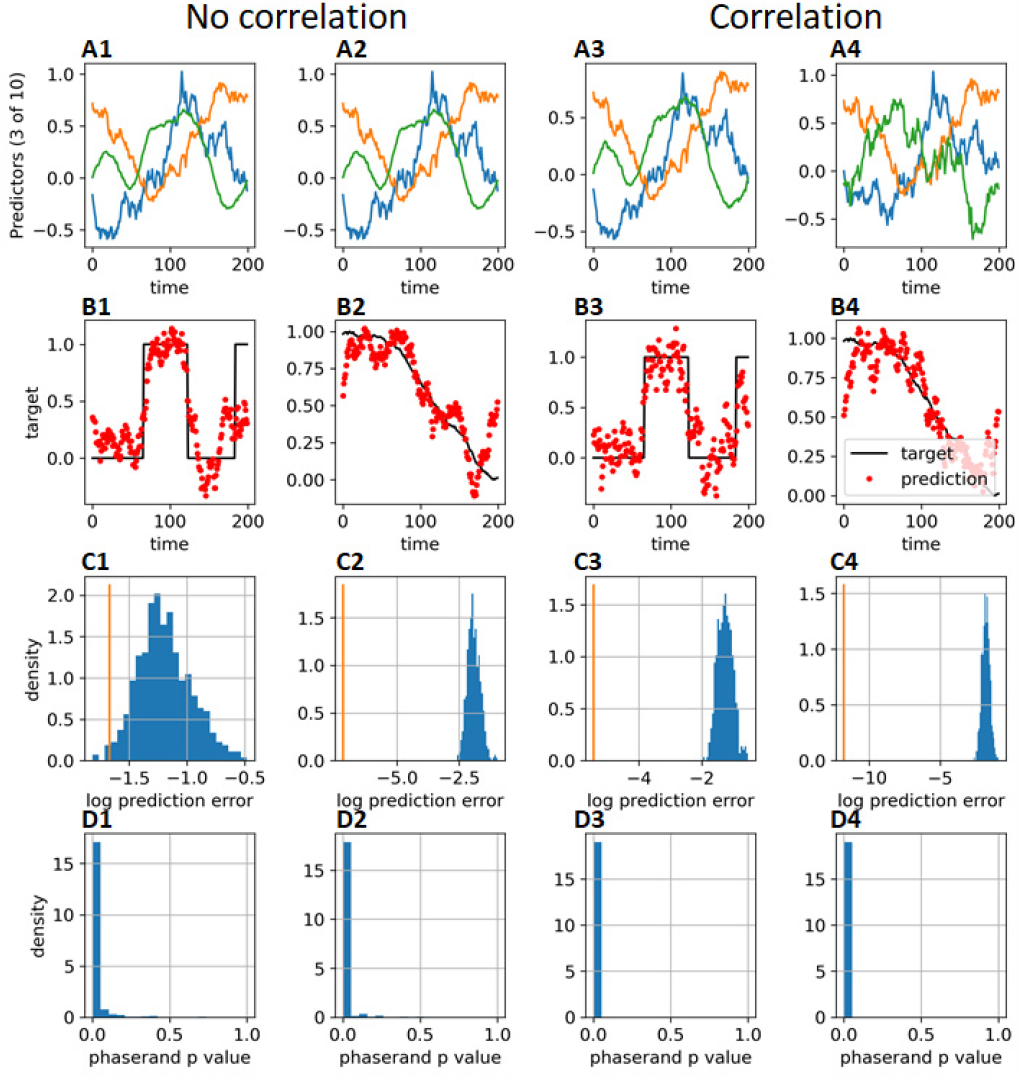
Phase randomization method. **A,B:** To create a null distribution, the neural data is randomly phase-shifted to create a surrogate timeseries with the same frequency content, and used to predict behavioral data. **C:** histogram of p-values obtained for the method, over 1000 simulations. Columns refer to the four scenarios of Figure 1.

An alternative method is wavelet randomization (Bullmore et al., 2001), which creates a null distribution by performing a wavelet transform on one of the timeseries, permuting the coefficients at each scale, and then inverse transforming. We found that this method performed better than Fourier randomization, but still gave substantially more false positives than linear shift (**Figure 7)**.

**Figure 7.**
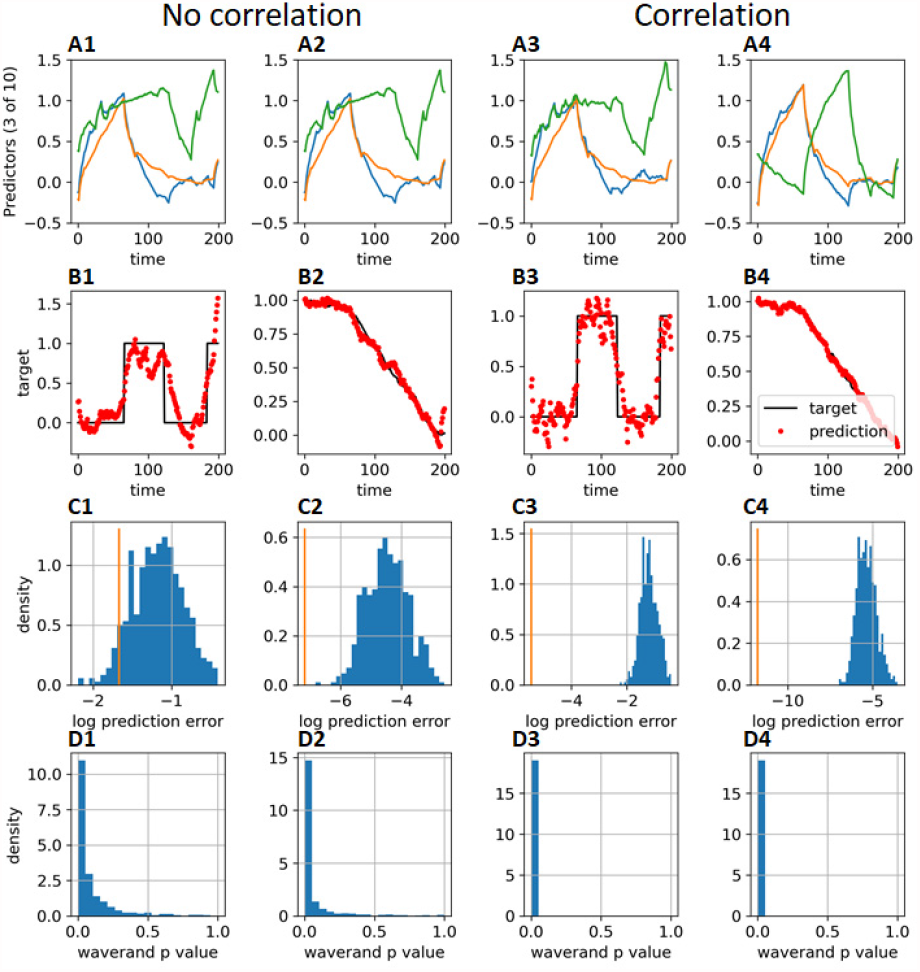
Wavelet randomization method. **A,B:** To create a null distribution, a Daubechies(4) wavelet transform is applied to the neural data, the wavelet coefficients are permuted within each scale, and inverse transformed create a surrogate timeseries with the same frequency content, and used to predict behavioral data. **C:** histogram of p-values obtained for the method, over 1000 simulations. Columns refer to the four scenarios of Figure 1.

## Cross validation

Cross validation does not solve nonsense correlations: slow autocorrelations mean that a predictor function learned on one part of the data will still be valid on another part of the data, even if these training and test sets are temporally segregated.

To demonstrate this, we applied 10-fold cross-validation to our four scenarios (**Figure 8)**. When the training and test sets consisted of random time points, performance was abysmal: test-set predictions of the behavioral variable were more accurate than predictions made without access to the simulated neural variables in 100% of simulations, even when the neural and behavioral variables were unrelated (**Figure 8A,B**). When training and test sets consisted of blocks of sequential trials, false-positives errors were less common but still occurred in 54% of simulations of the block behavioral variable and 92% of simulations of the continuous variable (**Figure8C,D**).

**Figure 8.**
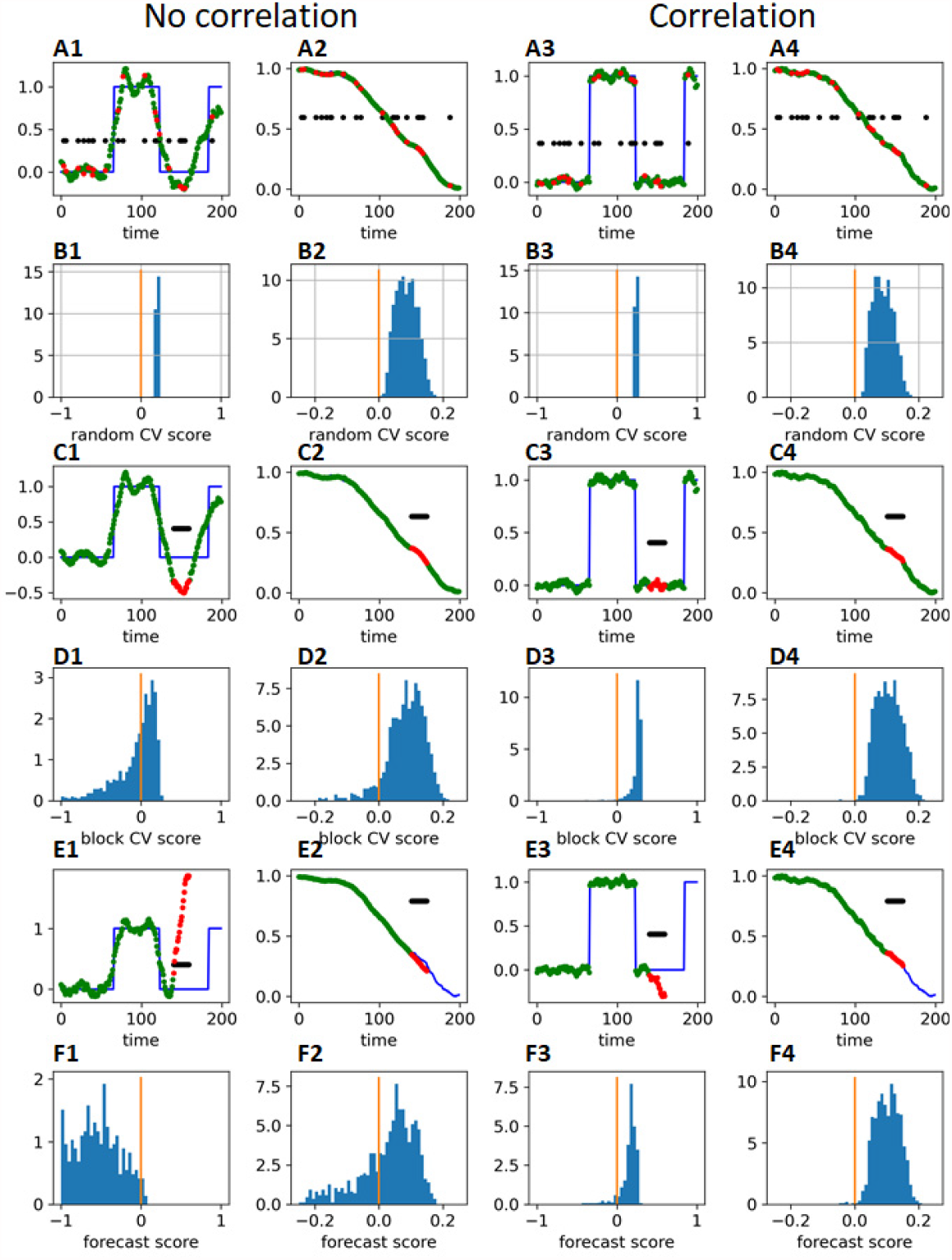
Cross-validation method. **A:** Trials were divided into 10 random sets, with 9 used to train a classifier (green dots) and the last to test the prediction (red dots). Black dots indicate a null prediction made without access to the neural variables (the training set mean). **B**: histogram of predictability (mean error of prediction without neural variables minus mean error with them), over 1000 simulated sessions. **C,D**: same analysis, with blocks chosen to be temporally continuous. **E,F:** forecasting method, where the training set (green) is strictly before the test set (red). Columns correspond to the four scenarios in Figure 1.

An alternative approach to time series cross-validation is forecasting (Tashman, 2000). In the approach, we predict the target timeseries in the *n*^*th*^ block using a predictor learned only from temporally prior blocks. As such predictions are extrapolation rather than an interpolation, one might expect false predictability to therefore be lower. This approach worked for the block variable, reducing the false positive rate to 1%; but it did not work for the continuous target, for which false-positives still occurred 61% of the time. (**Figure 8C,D**).

We conclude that cross-validation does not in general avoid nonsense correlations, although forecasting cross-validation can help in some circumstances. The use of cross-validation to avoid nonsense correlations must therefore be justified on a case-by-case basis.

## Auto-decorrelation

A commonly-suggested approach to eliminate nonsense correlations is to preprocess the data to remove correlations within a single timeseries (Haugh, 1976). If we could remove these autocorrelations, then standard statistical tests that assume independent samples could safely be applied to the auto-decorrelated timeseries.

The usual way to perform auto-decorrelation is with an autoregressive model: one predicts the value of **x**_*t*_ by linear regression from previous values **x**_*t*−*n*_ … **x**_*t*−1_, and performs all further analyses on the residual. Simpler approaches are to take the time derivative of each timeseries, or to detrend (for example by subtracting a best fit line).

While auto-decorrelation is in principle a solution to nonsense correlation, it comes with a major caveat: the auto-decorrelation algorithm must be extremely accurate. This cannot be guaranteed. For example, autoregressive models only exactly decorrelate linear timeseries (filtered white noise).

To evaluate auto-decorrelation, we fit a first-order autoregressive model to our simulated neural and behavioral variables, then applied a standard F-test to measure significance (**Figure 9)**. Because these timeseries are nonlinear, however, the autoregressive model did not fully decorrelate the data: slow trends were still observed in the neural data (**Figure 9A,B)** although smaller than prior to preprocessing. For the binary block variable, auto-decorrelation replaced the step functions with impulses at the start and end of each block. Even after autodecorrelation, the F-test produced inflated false positive rates, although this was less bad for the block variable (**Figure 9C**).

**Figure 9.**
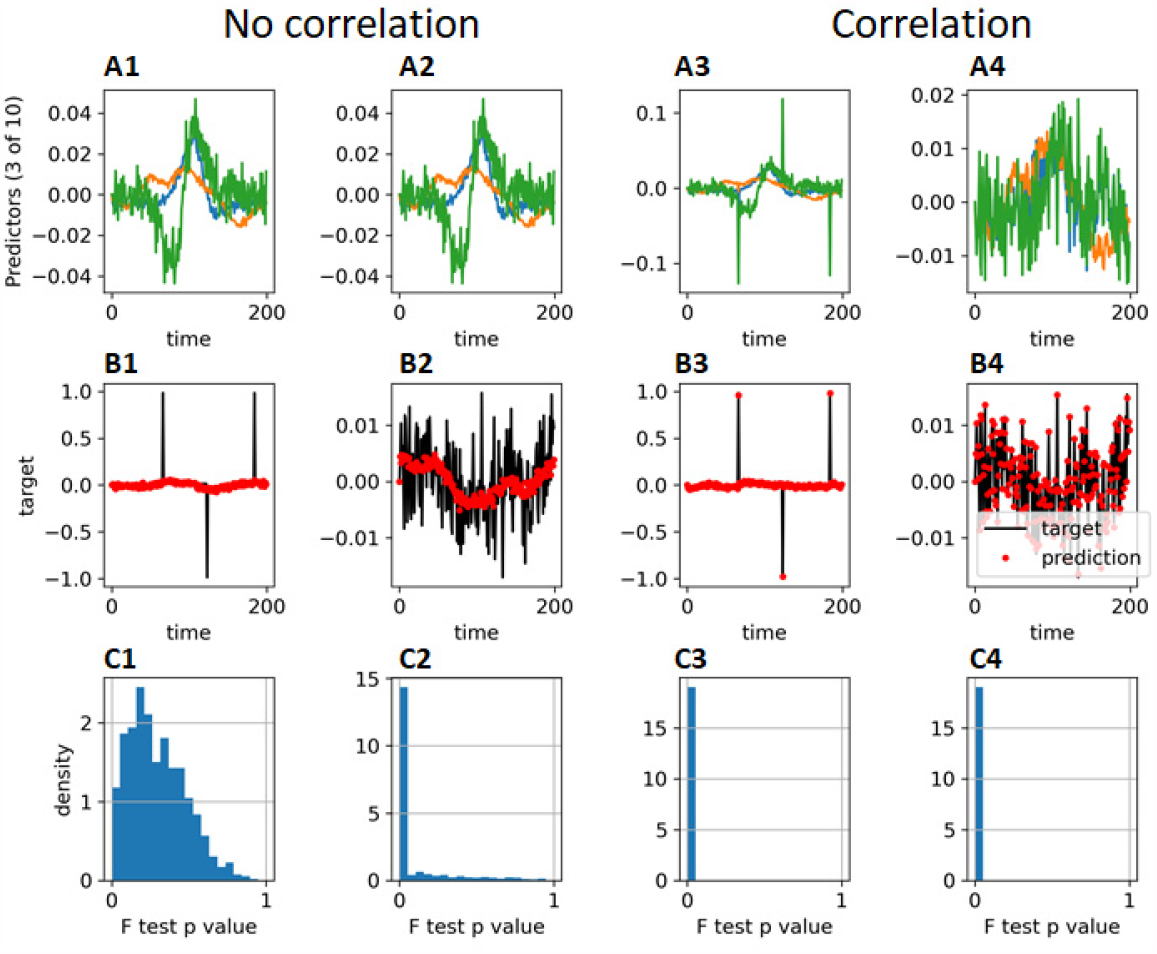
Auto-decorrelation. **A:** the simulated neural data was preprocessed by fitting a first-order autoregressive model to each timeseries, and calculating a residual at each timestep. Due to the nonlinear nature of these timeseries, this does not result in white noise. **B:** the behavioral variables were preprocessed the same way (black) and predicted from the preprocessed neural data (red dots) using multiple linear regression. **C:** histogram of F-test p-values measuring significance of the linear regression, over 1000 simulations.

Thus, it is not safe to apply statistical tests that assume independence even after auto-decorrelation, unless one has evidence that the auto-decorrelation method really works to high accuracy. In practice, this can be assessed by measuring how often the null hypothesis is rejected in synthetic data where there is known to be no genuine association. This approach has been used to validate auto-decorrelation in fMRI data (Afyouni et al., 2019; Cliff et al., 2021). Nevertheless, auto-decorrelation should be carefully tested before applying to neurophysiological data as they are substantially less linear than fMRI data. Linear timeseries are filtered independent noise; while fMRI timeseries look like filtered noise, neuronal unit recordings usually do not. For example, recorded cells can disappear halfway through an experiment due to technical issues such as electrode drift, which would not be corrected by a linear autoregressive model.

When auto-decorrelation does not fully decorrelate the data, it cannot be followed by statistical tests assuming independence of samples. Nevertheless, autodecorrelation can still be a useful tool used in conjunction with other approaches such as the linear shift method. Even if it only works partially, decorrelating the data cannot increase false positives found by the linear shift method, and may decrease them.

## More complex analyses: partial correlation

So far, we have discussed the simple case of detecting a correlation between two timeseries, such as neural activity and a behavioral variable. One often wants to ask more complex questions. For example, a subject’s choices differ between behavioral blocks. If neural activity correlates with behavioral block, is this correlation stronger than if the neuron encoded of choice alone? In other words, is there a partial correlation of neural activity and block, after accounting for the common correlate of choice? With autocorrelated timeseries, this is a much harder problem than simply testing for independence.

We must first carefully define what we mean by partial correlation for autocorrelated timeseries. Suppose we are predicting a vector timeseries **Y** from another vector timeseries **X**, and we also measure a third confounder timeseries **Z** which is not independent of **X**. For example, **Y** might contain population activity on each trial, **X** might contain the behavioral block on each trial, and **Z** the choice on each trial. Because the subject learns from previous trials, **Z** and **X** are not independent, but we do not have an equation accurately modeling their relationship.

We would like to test a null hypothesis that **Y** has no correlation with **X** beyond that explained by dependence on **Z**. This can be formalized as a model **Y** = **ZW** + **E**, where **W** is a non-random but unknown weight matrix, and **E** is independent of **X** (Harris, 2021). As before, this means independence of the entire histories over repeats of the experiment: ℙ(*e*_*ct*_, *x*_*du*_) = ℙ(*e*_*ct*_)ℙ(*x*_*du*_), for any cell *c*, any dimension *d* of the confounding variable, and any pair of times *t* and *u*.

Even if we know the probability distribution of ℙ(**X**), we cannot use the pseudosession method to test independence of **E** and **X**, since we do not measure **E** directly and we do not know ℙ(**Z**|**X**). We also cannot use session permutation, since no one of our three observed series **X, Y, Z** is independent of the other two under the null.

We recently described an approximate test for partial correlations between autocorrelated timeseries null where one has observed multiple experiments (Harris, 2021). Let ***X***_*i*_, ***Y***_*i*_ and ***Z***_*i*_ be the timeseries observed on experiment *i*, and let ***P***_*i,j*_ be the *T*× *T* projection matrix orthogonal to all columns of both ***Z***_*i*_ and ***Z***_*j*_. Given a user-supplied measure *ρ*(***X***; ***Y***) of association between two timeseries (for example Pearson correlation), we define the statistics

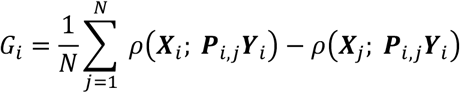

Under the null, the *G*_*i*_ are mutually independent and their sum has mean 0 (Harris, 2021). If their distributions are close to symmetric (which can be checked with histograms or QQ plots), the null can be tested using a t-test that the mean of *G*_i_ is 0 (Cressie, 1980; Efron, 1969).

## Implications for experimental design

This survey of methods for establishing genuine correlations between neural and behavioral timeseries yields a familiar lesson for experimental design: whenever possible, use a randomized experiment.

The power of randomized experiments to enable statistical analysis has long been recognized (Fisher, 1935). Of the methods described above, the one that is most reliable, powerful, and accurate is the pseudosession method, which can only be applied when one of the timeseries to be correlated is randomly controlled by the experimenter. Thus, whenever possible, experiments should be designed with randomized covariates. To test if neural population activity differs between behavioral blocks, the timing of these blocks should be randomized between sessions. To test if neural activity correlates with running, the best experimental design would be one that requires the subject to run at random times controlled by the experimental apparatus.

## Summary

We have reviewed and simulated methods for detecting correlations between neural timeseries. Statistical tests that assume independence between timepoints result in “nonsense correlations” of erroneous statistical significance, due to autocorrelations within timeseries. The most reliable method for detecting genuine correlations, the pseudosession method, requires that one of the timeseries be randomly generated by the experimenter. The session permutation method requires the experiment be replicated at least 5 times, and can provide reliable results although correlations could reflect a common effect of session-to-session variability. The conservative linear shift method works if one of the series is stationary. Other methods reduce but do not eliminate the risk of false positive errors. If possible, experiments should be designed so that time series of interest are randomized.

## Methods

To generate a firing rate sequence, we added together a random number of logistic sigmoid functions. The center times *t*_1_,… *t*_*n*_ of these functions were drawn from a homogeneous Poisson process of rate 1/100, so the mean number of sigmoids in T=200 trials was 2; their widths were always 10, and directions *σ*_*i*_ were random signs ±1 with equal probability. A pink noise sequence *p*_*t*_ was added, generated by passing white Gaussian noise through an IIR filter with parameters 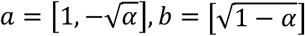, where *α* = *e*^−2/*τ*^ and *τ* = 5000. The final sequence was

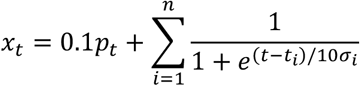

To simulate behavioral binary blocks (column 1 of the figures), we generated alternating blocks of 0s and 1s, of lengths independently uniformly distributed between 50 and 70; the first 0 block always began at the first sample. To make a stationary block sequence (Figure 5), we generated a longer sequence (1000 blocks) and started it at a random time. To simulate continuous behavioral signal (column 2 of the figures), we generated another sequence from from the same distribution as the neural activity.

To simulate the case where the neurons encoded information about the behavioral variables (columns 3 and 4), the behavioral signal was added to each neuron’s activity, with weight drawn from a Gaussian distribution of mean 0, SD 0.1.

Finally, each neuron’s activity timeseries scaled between 0 and 1, although this will not have affected the linear regressions.

Code to perform the simulations is available at https://github.com/kdharris101/nonsense-correlations/ and can be run online at https://colab.research.google.com/github/kdharris101/nonsense-correlations/blob/main/nonsense.ipynb

## Acknowledgements

I thank Sylvia Schroeder, Kevin Miller, Anna Lebedeva, Peter Dayan, Tim Behrens, Steve Smith, and Matteo Carandini for conversations and comments on the manuscript. This work was supported by the Wellcome Trust (205093) and European Research Council (694401).

